# Distinct latitudinal patterns of shifting spring phenology across the Appalachian Trail Corridor

**DOI:** 10.1101/2023.12.11.571108

**Authors:** Jordon Tourville, Georgia Murray, Sarah Nelson

**Author notes:** **Corresponding Author:** Jordon Tourville. **Open Research Statement:** Data are provided as private-for-peer review in GitHub via the following link: [https://github.com/jtourvi/AT_Phenology.git].

## Abstract

Warming associated with climate change will likely continue to advance the onset of spring phenology for many forest plants across the eastern United States. Understory forbs and spring ephemerals which fix a disproportionate amount of carbon during spring may be negatively affected by earlier canopy closure (i.e., phenological windows), however, information on the spatial patterns of phenological change for these communities is still lacking. To assess the potential for changes in spring phenological windows we synthesized observations from the Appalachian Mountain Club’s (AMC) Mountain Watch (MW) project, the National Phenology Network (NPN), and AMC’s iNaturalist projects between 2004 and 2022 (n = 118,250) across the length of the Appalachian Trail (AT) Corridor (34°N-46°N latitude). We used hierarchical Bayesian modeling to examine the sensitivity of day of year of flowering and leaf-out for 11 understory species and 14 canopy tree species to mean spring temperature (April-June). We conducted analyses across the AT Corridor, partitioned by regions of 4° latitude (South, Mid-Atlantic, and North). Spring phenologies for both understory plants and canopy trees advanced with warming (∼6 days/°C and ∼3 days/°C, respectively). However, sensitivity of each group varied by latitude, with phenology of trees and understory plants advancing to a greater degree in the mid-Atlantic region (∼10 days/°C) than the southern or northern regions (∼5 days/°C). While we find evidence that phenological windows remain stable in southern and mid-Atlantic portions of the AT, we observed an expansion of the spring phenological window in the north where there was greater understory temperature sensitivity compared to trees (∼1.6 days/°C). Our analyses indicate differential sensitivity of forest plant phenology to potential warming across a large latitudinal gradient in the eastern United States. Further, evidence for a temperature-driven expansion of the spring phenological window suggests a potential beneficial effect for understory plants, although phenological mismatch with potential pollinators is possible. Using various extensive citizen-science derived datasets allows us to synthesize regional- and continental-scale data to explore spatial and temporal trends in spring phenology related to warming. Such data can help to standardize approaches in phenological research and its application to forest climate resiliency.

## Introduction

### Spring phenology and changing climate

Phenology represents the timing of critical life events for plants, both over their entire lifespan and on an annual cycle (see Table 1 for definitions of important terms used throughout this article; Cleland et al., 2007; Piao et al., 2019). The timing of events, such as flowering, bud-break, fruiting, and leaf senescence, and the synchronicity between these events and important climate and biotic interactions (pollination, seed dispersal, etc.) can dictate the performance (i.e., fitness) of plant individuals or even entire populations (Aerts et al., 2006). For example, spring flowering timing of understory ephemerals in eastern North America is beneficial for plants when synchronous with the peak activity of generalist insect pollinators, and when occurring after late-season frosts (Neufeld and Young, 2003; Inouye, 2008; Rafferty and Ives, 2011; Ettinger et al., 2018). In the former case, plant reproduction is enhanced through specific flowering phenology, and in the latter, plant growth and survival are improved when flowering avoids freezing temperatures. Phenological events are controlled tightly by physiological mechanisms which rely on environmental cues like temperature and photoperiod (Neufeld and Young, 2003; Gilliam, 2007; Wang et al., 2020; Moon et al., 2021).

**Table 1:** Definitions of important terms used throughout the study.

| Term | Definition |
| --- | --- |
| Phenology | Timing of recurring plant life stages and their relationships with weather and climate |
| Spring ephemeral | Perennial woodland wildflowers which develop aboveground early each spring, bloom, produce seed, and senesce during a brief window of time |
| Leaf-out | Leaf expansion following leaf budburst during spring months |
| Canopy closure | Full overlap of expanded tree canopy leaves which fully shades the forest understory |
| Spring phenological window | Brief period of high light conditions below the forest canopy starting with understory leaf-out and ending with full canopy closure |
| Phenological mismatch | Interacting species change the timing of regularly repeated phases in their life cycles at different rates |
| Hopkins' Bioclimatic Law | Hypothesized phenological pattern where a 4-day shift in phenology is predicted for a change of every 1° latitude north, 5° longitude west, and 120 m increase in elevation |
| Appalachian Trail Corridor (AT) | Lands immediately surrounding the Appalachian Trail that follow the Appalachian Mountain chain from Georgia to Maine (USA) |
| iNaturalist | An online platform that allows sharing of biodiversity observations and creates research-quality citizen science data |
| Appalachian Mountain Club (AMC) | An environmental non-profit organization dedicated to conservation, education, and recreation of lands within the AT Corridor |
| National Phenology Network (NPN) | An organization established in 2007 to collect, store, and share phenology data and information |
| Mountain Watch Project (MW) | An AMC project established in 2004 to monitor phenological change in high elevation environments within the AT Corridor |
| Phenophase | An observable stage in the annual life cycle of a plant that can be defined by a start and end point |
| Day of Year (DOY) | Julian date for which a particular phenophase is observed for an individual plant |
| Intensity values | Ordinal categories for a particular phenophase |
| Plant performance | Integrated measure of plant fitness which incorporates survival, growth, and reproductive success |
| Forest climate resilience | A measure of a forest's adaptability to a range of climate stresses which reflects the functional integrity of the ecosystem |

In seasonal temperate forests, understory plants, which include spring ephemerals, perennial forbs, shrubs, and tree seedlings, spend the majority of the growing season in low-light conditions (Gilliam, 2007). To compensate, many of these species undergo bud break and leaf expansion (referred to as leaf-out) prior to full canopy closure in the spring (Neufeld and Young. 2003; Lee and Ibanez, 2021a). In an environment with such high light-availability, understory species can fix the majority of their annual carbon budget in this time period, in some cases up to 80-90% (Kudo et al., 2008; Augspurger and Salk, 2017; Heberling et al., 2018; Lee and Ibanez, 2021b). Thus, the brief period of time (referred to as the phenological window) between understory leaf-out and canopy leaf-out is essential for understory plant performance (Heberling et al., 2019).

Given the tight coupling of spring phenology and climate, warming caused by climate change could alter the dynamics of the phenological window in several ways (Figure 1). Importantly, current evidence suggests that different plant functional groups (understory herbaceous plants and some shrubs *vs.* canopy trees) respond to different sets of environmental cues, meaning that spring phenology of each could be altered asynchronously given accelerating climate change and other global change drivers (Richardson and O’Keefe, 2009; Lee et al., 2022; Alecrim et al., 2022; Miller et al., 2023). For instance, canopy trees may be more responsive to air temperature (directly influenced by warming), while understory species may be relatively more sensitive to soil temperature and snow depth (Zohner et al., 2016; Jánosi et al., 2020). In a situation where canopy closure advances with air temperature increases at a greater rate than understory leaf-out, understory species may suffer from lower photosynthetic rates leading to reduced carbon gain, which may in turn have ecosystem-level consequences (Beard et al., 2019; Heberling et al., 2019). Monitoring the phenological response of both forest canopy trees and understory plants is essential for understanding the risks posed by climate change in these systems.

**Figure 1:**
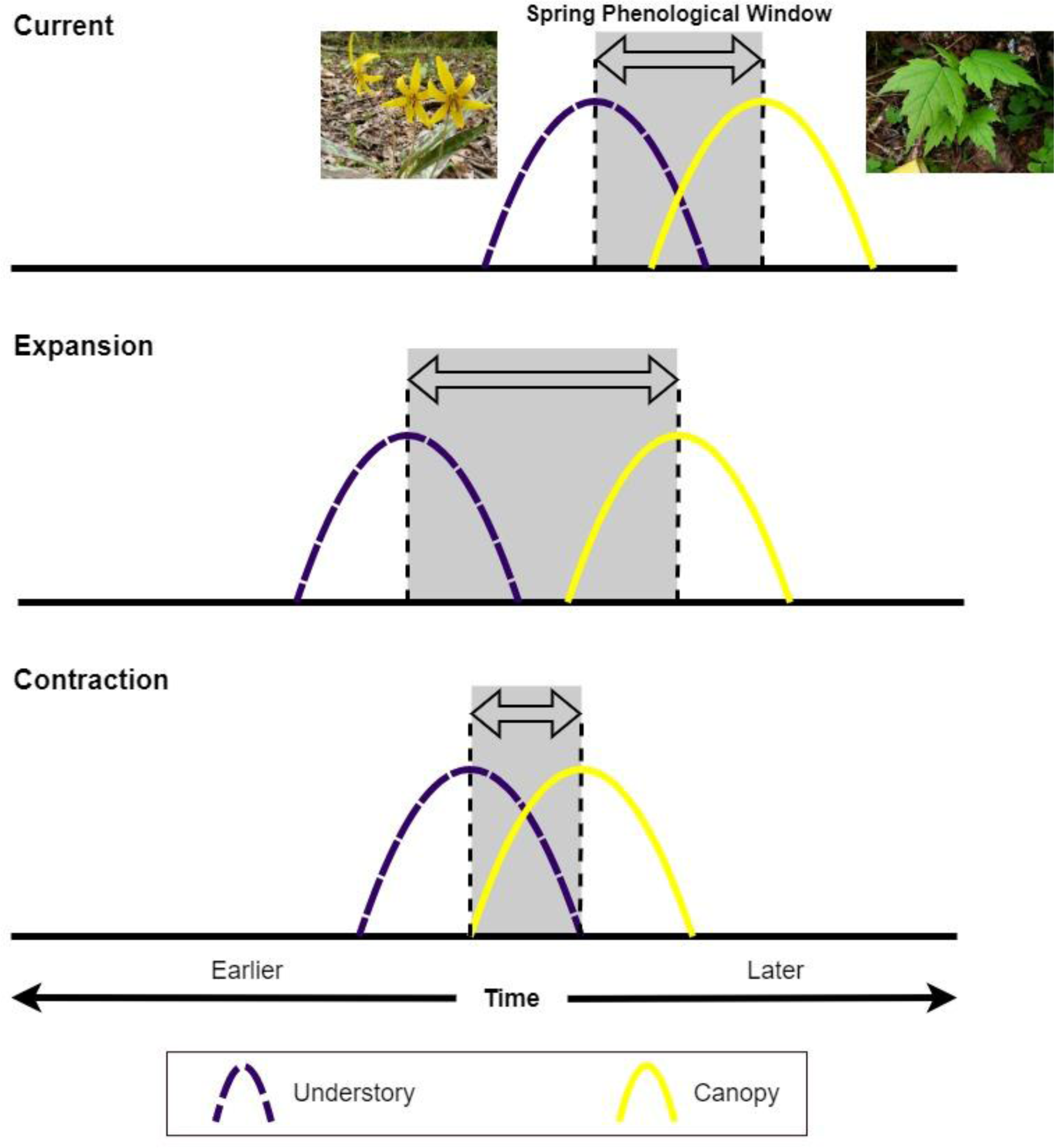
Conceptual diagram depicting different scenarios for the shifting spring phenological window under warming. Under the current scenario, both co-occurring understory (purple dashed curve) and canopy (yellow solid curve) species advance their phenologies at the same rate, keeping the window (shaded area) the same length through time. In an expansion scenario, understory plants advance their phenologies faster than canopy trees, leading to a greater period of high-light conditions for these species. Under the contraction scenario, canopy trees shift faster than understory plants with warming, reducing the period of high-light conditions for understory species and possibly leading to reduced annual carbon acquisition. Credit for all photos: Jordon Tourville.

### Contrasting patterns of phenological shifts

Recent findings in eastern North America have illustrated greater advances of spring phenology for canopy trees compared to spring-blooming understory herbaceous species over a 160-year period, leading to a potential future understory carbon budget loss of 12-26% from increased shading (Heberling et al., 2019). Another study estimating spring phenology from herbarium records came to similar conclusions for forests in eastern North America, although the spring phenological window remained stable with warming in European and East Asian forests (Lee et al., 2022). However, both Ge et al. (2015) and Alecrim et al., (2022) reached the opposite conclusion, finding an expansion of the spring phenological window within forests in China and the eastern United states, respectively, which, barring other phenological mismatches (i.e., with pollinators), could be a net positive for understory species. Thus, based on conflicting evidence there is no consensus on whether trees or understory plants are advancing their phenology more strongly in response to climate change.

These recent conflicting findings may be the result of challenges stemming from the high degree of environmental, geographic, genetic, and methodological variation encountered. For example, higher-latitude regions are warming faster than others (particularly in spring and winter), which could suggest greater magnitudes of phenological change over time in those locations if species’ phenologies are responding principally to temperature (Rice et al., 2018; Montgomery et al., 2020). However, population-level intraspecific variation in phenological temperature sensitivity may serve to blunt these responses (McDonough MacKenzie et al., 2018; 2019). Additionally, environmental variables both at a single site and across a wide geographic range, such as precipitation, elevation, and edaphic factors, could also affect how species track a changing climate (Du et al., 2020; Alecrim et al., 2022). This is particularly true given well-established geographical patterns, such as Hopkins’ Bioclimatic Law which hypothesizes a 4-day shift in phenological events for every 1° latitude north, 5° longitude west, and 120 m increase in elevation (Hopkins, 1920). Additionally, the diversity of phenological data used to estimate the spring phenological window (i.e., wildflower leaf-out *vs.* flowering, herbaria records *vs.* direct observation *vs.* experimental manipulation) could lead to different conclusions (Wolkovich et al., 2012; Heberling et al., 2019; Alecrim et al., 2022; Lee et al., 2022). To help resolve these discrepancies we need spatially and temporally extensive, multi-sourced phenological datasets comprised of different functional groups that represent variation in climate and topography across a large geographic area.

### Appalachian Trail Mega-transect and citizen science

The Appalachian Trail (AT) Corridor and its surrounding 250,000 acres of federally protected lands form the AT Mega-transect (Cohn, 2008). This corridor harbors rare, threatened, and endangered species, encompasses important water resources, and shelters a high diversity of wildlife (Cohn, 2008). The AT’s north-south alignment across 14 states represents a lengthy (12°) latitudinal gradient within the eastern United States and offers an ideal setting for collecting relevant phenological data on a continental scale (Wang, 2020a; Wang, 2020b). Threats to the environment of the AT—from encroaching development, acid rain and air pollution, invasive species, polluted water, and climate change—represent threats to the health of everyone downwind and downstream of the AT, roughly one-third of the U.S. population (McKinley et al., 2019; Burns et al., 2020). Thus, the AT Corridor thus serves as an excellent monitoring nexus for environmental conditions that directly affect more than 120 million Americans (McKinley et al., 2019). The dense population and abundant recreational opportunities near and within the AT Mega-transect also allow for ample community research engagement.

Community science, or citizen science, is the practice of engaging the community to participate and collaborate in scientific research (Wandersman, 2003; Tebes, 2005; Cooper et al., 2021). This method of data collection is a useful tool to expand spatial coverage of monitoring projects that would be otherwise hindered by funding and resources. Additionally, community scientists can help directly support conservation efforts, and build meaningful connections to their community and natural environment (Bonney et al. 2016). With long-term phenology monitoring supplemented with thousands of community scientist observations through platforms like iNaturalist (Table 1), changes and shifts in phenological responses to warming can be identified along the AT Corridor (Nugent, 2018; Soroye et al., 2022). Observing plant phenology along the AT may allow us to better account for the high spatial and environmental variability common in studies investigating shifting phenology with climate.

### Study questions

Given the uncertainty around the direction and magnitude of changes to the spring phenological window under a changing climate, as well as the need to understand the effects of warming on forests within the AT Corridor, it is imperative that we use spatially extensive phenological datasets. Using data from the Appalachian Mountain Club’s (AMC) Mountain Watch (MW) Project, National Phenology Network (NPN), and iNaturalist, we first determined what climate or landscape factors are relevant drivers of spring phenology for our focal species (14 canopy tree and 11 understory species). Using this information, we asked, (Q1) is spring phenology of canopy trees and understory forbs and shrubs advancing with warming, (Q2) and if so, are there differences in phenological sensitivity to temperature between these groups (i.e., causing a phenological mismatch)? We also asked (Q3) are there differences in the magnitude of phenological shifts across the length of the AT corridor (∼12° latitude) and by individual taxa?

## Methods

### Study region

Our study area includes all temperate broadleaf forests within HUC10 (U.S. Geological Survey hydrological units, https://irma.nps.gov/DataStore/Reference/Profile/2184124) watersheds that intersect the Appalachian Trail and surrounding Corridor. Temperate forests of eastern North America are characterized by seasonality, with high light penetrating to the forest floor in the shoulder seasons (early spring and late fall), and low light under closed tree canopy during warmer months in the growing season (down to 1-5% canopy openness, Beeles et al., 2022). Northern hardwood communities comprising sugar maple (*Acer saccharum*) and American beech (*Fagus grandifolia*) dominate northern AT regions, while oak-hickory forests (*Quercus sp.* and *Carya sp.*) are common along the southern AT within lower elevations (Tourville et al., 2022; Janowiak et al., 2018). At higher elevations, evergreen montane spruce-fir forests proliferate, although these do not display the same seasonality as lower-elevation broadleaf forests. In the understory, herbaceous forbs, including spring ephemerals such as *Dicentra sp. and Erythronium sp.*, shrubs such *as Viburnum sp.*, and tree seedlings of overhead canopy species are common (Heberling et al., 2019; Tourville et al., 2022).

The AT is the longest footpath in the world (∼2,190 miles), traveling through 14 U.S. states from its southern terminus, Springer Mountain, Georgia, to its northern terminus, Katahdin in Maine (see Figure 2). Over 3 million people visit the trail each year, making it an ideal monitoring corridor for community science efforts, and where shifts in phenology can be recorded over large geographic extents (Cohn, 2008). From Southern Appalachian grassy balds, to the alpine zones of the Northeast, the AT is also home to diverse flora that may be influenced by climate change. Higher latitudes in the Appalachians are warming and experiencing longer growing seasons but elevational differences are mixed (Kimball et al., 2014; Janowiak et al., 2018; Murray et al. 2021). While data from many other global montane sites have demonstrated elevation-dependent warming (Pepin et al. 2022) this has not been the consistent pattern in the northern Appalachians (Murray et al. 2021).

**Figure 2:**
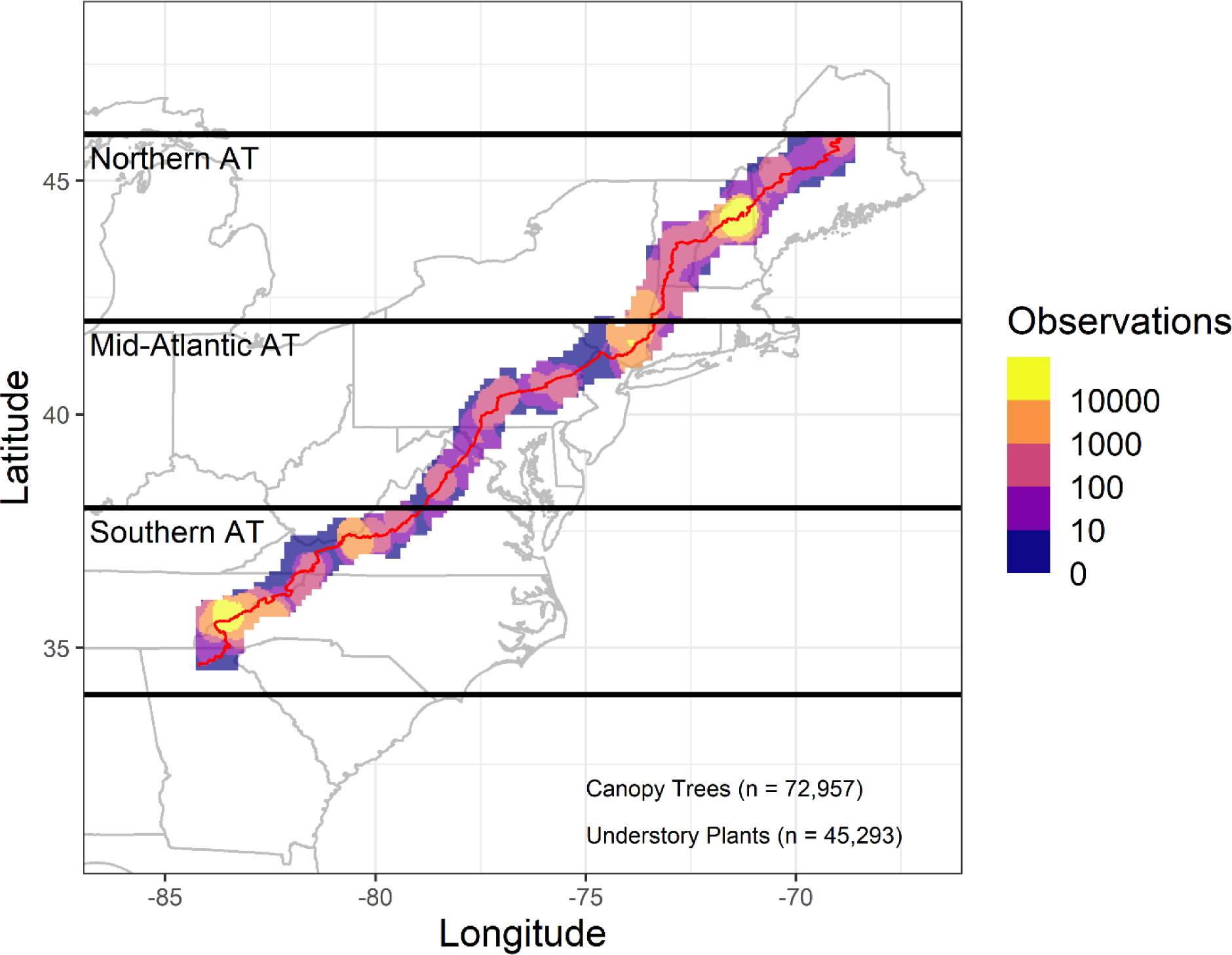
Point density estimates of canopy tree and understory plant spring phenology observations analyzed in this study across all HUC10 watersheds intersecting the AT between 2004-2022 (n = 118,250). The red line indicates the Appalachian Trail. Note that the scale is logarithmic.

### Data collection

In 2004, AMC began monitoring reproductive plant phenology events focused on flowering for alpine species, and later expanded to include woodland species (trees, shrubs, and forbs) and other phenophases such as leaf-out in northeastern mountains in the United States. Initially designated the Mountain Watch (MW) project, this effort enlisted organizational staff, partner organizations, and volunteers to gather phenology data on paper data sheets. The MW Project has since evolved to utilize the National Phenology Network’s (NPN) protocol and currently collects data in two primary ways: through (*i*) the establishment of permanent plots in the White Mountains of New Hampshire, and (*ii*) by using phone applications (apps) and smartphones to enhance monitoring practices. In some cases, partner organizations have also set up permanent plots and similarly evolved to use the NPN protocol. In recent years, monitoring through community science has expanded from the Northeast to the entire AT Corridor using the platform iNaturalist.

iNaturalist is a free smartphone app with currently 2.5 million active users and nearly 70 million observations (Barve et al., 2020; Callaghan et al., 2022). Users can upload photo observations, provide a species ID, or receive one based on the program’s algorithm or a community of online naturalists. Observations are made research grade once there are two corresponding species identifications. The iNaturalist geotagged images also reduce location errors and eliminate the past challenge of inaccurate species ID from novice observers (McDonough MacKenzie et al. 2017; McDonough MacKenzie et al. 2020). iNaturalist serves as a supplement to permanent plots as NPN plots require consistent attention from skilled naturalists while having limited spatial distribution. Importantly, iNaturalist observations can be used to fill gaps between monitoring plots and expand spatial and temporal data coverage. Researchers can also create projects on iNaturalist to capture observations of a specific species or geographical range.

AMC’s iNaturalist phenology projects incorporate NPN’s protocol as observation fields to identify the phenophase of a plant observation. Staff project curators and managers have the task of adding observations of target species within the AT Corridor to the project if they have not been uploaded to the project by the observer. AMC’s iNaturalist project, *Flowers and Fauna along the Appalachian Trail Corridor*, began in 2018 and with continued dedicated funding has grown to now include >40,000 phenological observations (see Supplemental Figure S1). AMC’s ultimate goal is to establish a long-term dataset that can be expanded and analyzed year after year to infer changes in plant phenological responses to changing climate along the full AT Corridor.

### Data Preparation

We synthesized and collated phenological observations from three sources: the AMC’s MW Project, the NPN online data portal, and the AMC’s iNaturalist projects (>2 million observations). Observations ranged from 2004 to the end of 2022 and represented multiple phenophases (leaves, flowering, fruiting, senescence, etc.), and plant species (understory woodland, canopy trees, and alpine species). As we were only interested in spring leaf and flower phenology, we removed observations recorded of other phenophases and from other seasons from the dataset. Additionally, we used only positive observations of phenology in subsequent analysis (i.e., only records where the phenophase was actually observed). Since we were only comparing spring phenology of understory and tree species in temperate broadleaf forests, we removed alpine species observations from the dataset. Further, we removed understory herbaceous species that either flower later in the growing season (i.e., after canopy closure), or had fewer than 100 observations. We only kept tree species that had the potential to maintain a dominant position in the forest canopy and had greater than 100 observations. We used ArcPro v3.1 (ESRI, 2022) to create a watershed delineation buffer using USGS HUC10 watersheds around the AT. We included only records within our HUC10 AT buffer in our analysis (in the eastern United States between approximately 34-46°N latitude).

All observations were highlighted for either day of year (DOY) of leaf-out (tree species and understory species, DOY_leaf_) or DOY of open flowering (understory species, DOY_flower_), consistent with previous studies comparing phenology across forest strata (Heberling et al., 2019; Lee et al., 2022). NPN defines leaf-out as one or more individual leaves unfolded, meaning the entire length of the leaf has emerged from the bud (NPN, 2023). Flowering is defined as when one or more flowers are open so that reproductive parts are visible (NPN, 2023). Our approach assumes that herbaceous species’ flowering and leaf-out timing is tightly correlated for these species (see Lee et al., 2022). Understory flowering may be a better choice of response over leaf-out because intensity values, or ordinal categories for each phenophase (e.g., <5%, 5-25%, 25-50%, 50-75%, 75-95%, >95%), are measured and associated with understory flowering but not understory leaf-out. Like canopy tree DOY_leaf_ which also have associated intensity values, DOY_flower_ can be viewed as a distribution. Understory plants with or without leaves as a binary measure makes it difficult to truly assess true leaf-out timing.

We focused on common and ubiquitous species (found across the majority of the AT Corridor, minimum >4° latitude) that flowered and leafed out at approximately the same time in early spring. We subsequently removed observations where either leaf-out or flowering occurred after DOY 200, as these were either likely in error or a second flowering which occurs in some species under the right conditions. We also only included tree observations where budburst or leaf expansion intensity values were 75-95%, indicating nearly total leaf-out. In total, after accounting for observations that were excluded from our original search, we collected data for a total of 25 species (14 tree species and 11 understory forb and shrub species) consisting of 118,250 individual observations across the entire AT Corridor (Figures 2, 3). In order to examine latitudinal differences in spring phenology, observations were partitioned among three 4° latitudinal bands: Southern AT (34-38°N), mid-Atlantic (38-42°N), and Northern AT (42-46°N) (Figure 3).

**Figure 3:**
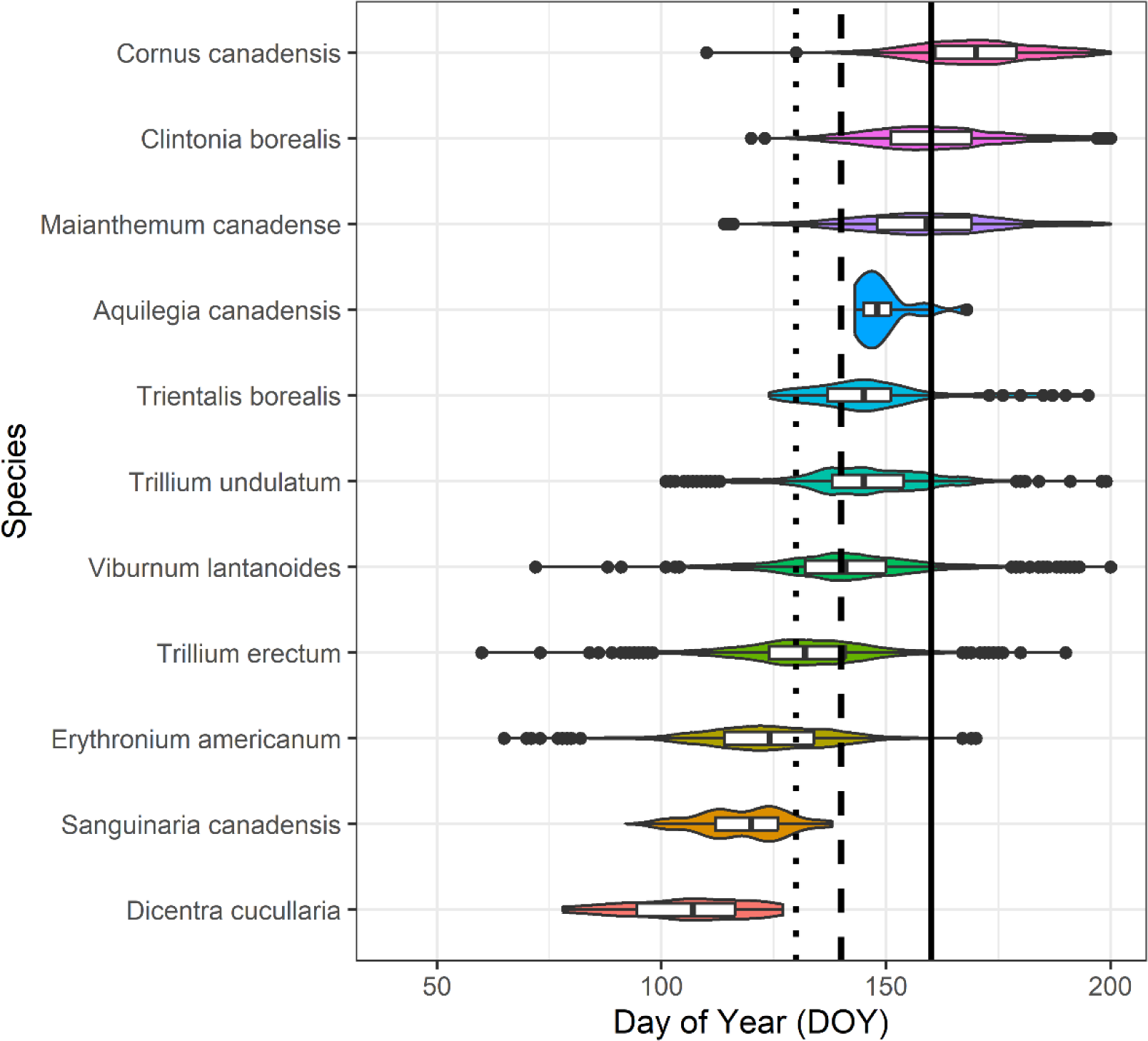
Density and distribution of understory flowering observations for each of the 11 species examined in this study. The observations shown are only from the entire AT Corridor (34-46° N latitude). The solid line displays the median canopy closure date (from observed citizen-science data) between 2004-2022 for the northern AT (DOY = 160, June 9^th^), while the dashed and dotted lines show median canopy closure date for the mid-Atlantic (DOY = 140, May 20^th^) and southern (DOY = 130, May 10^th^) AT regions, respectively. Asterisks (*) indicate species definitively identified as spring ephemerals. Boxplots show median values for DOY_flower_ (with 25% and 75% quantiles).

Since all observations were geolocated (we only kept observations with < 250 m accuracy error), we were able to extract potentially relevant landscape and climatological data for each record. DAYMET 1 km gridded climate data were extracted for the year an individual observation took place, which included mean, maximum, and minimum air temperatures (°C), solar radiation (W/m^2^), snow water equivalent (SWE, mm), vapor pressure deficit (Vpd, kPa), and total precipitation (mm) (Daymet: Daily Surface Weather Data on a 1-km Grid for North America, Version 4 R1 https://doi.org/10.3334/ORNLDAAC/2129). Daily values were extracted for the entire year which the observation was recorded allowing us to calculate daily, monthly, seasonal, and annual mean values for each climate variable. Accumulated growing degree days (AGDD) were also derived from DAYMET temperature data using Equation 1 (Gavin et al., 2008; Wason and Dovciak, 2017; Tourville et al., 2022):

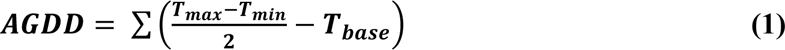

Where *AGDD* is the accumulated maximum value of growing degree days for spring only, *T_max_* and *T_min_* are daily maximum and minimum temperatures, and *T_base_* is a constant (4°C). Elevation at each observation location was extracted from a national 10 m digital elevation model (DEM).

### Data Analysis

In order to determine which candidate geographic and climate variables (Table 2) influence spring phenology for both understory and canopy species and warranted inclusion in subsequent models, a preliminary exploratory multiple regression analysis was undertaken using DOY_leaf_ and DOY_flower_ as a response. Our main candidate variables included mean spring temperature (April-June, AMJ), mean previous winter temperature (DJF), spring AGDD, seasonal SWE (proxy for snow depth, DJF) latitude, and elevation (see Table 2 for all possible candidate variables). Spring temperature was calculated as the average of the April, May, and June daily temperatures for the year and the location associated with each phenology record. April-June temperatures explained more variation in DOY than other spring windows (i.e., March-May, or individual months). All variables were scaled and centered and global models with all candidate variables were examined. For both understory and canopy species, standardized regression coefficients for mean spring temperature (negative interaction), elevation and latitude (both positive interactions) were significant predictors of DOY (Supplemental Figure S2). Regression models with spring temperature, latitude, and elevation explained more or similar variation (marginal R^2^) in DOY to more complex models, thus, we used these three variables in subsequent modeling of changes to the spring phenological window (see below).

**Table 2:** Abbreviations, definitions, and the data source of independent variables (by category) used for exploratory analyses for predicting DOY of tree leaf-out and understory flowering across the AT Corridor. Elevation, latitude and longitude were derived from geolocation data of individual observations and regional 2 m digital elevation models (DEM). Climate values were extracted from DAYMET 1-km gridded datasets. AGDD was derived from extracted DAYMET temperature data.

| Category | Variable | Abbreviation | Units | Description | Source |
| --- | --- | --- | --- | --- | --- |
| Geography | Elevation | Elev | m | altitude above sea level | AMC |
|  | Latitude | Lat | ° | degrees latitude | AMC |
|  | Longitude | Long | ° | degrees longitude | AMC |
| Climate | Spring maximum temperature (AMJ) | Spring Tmax | °C | maximum spring temperature for each observation year | DAYMET |
|  | Spring minimum temperature (AMJ) | Spring Tmin | °C | minimum spring temperature for each observation year | DAYMET |
|  | Spring mean temperature (AMJ) | Spring Tmean | °C | mean spring temperature for each observation year | DAYMET |
|  | Accumulated solar radiation | SR | W/m<sup>2</sup> | sum of incoming solar radiation for each observation year | DAYMET |
|  | Maximum snow water equivalent (DJF) | SWE | mm | maximum snow water equivalent for observation winter | DAYMET |
|  | Accumulated growing degree days | AGDD |  | sum of daily temperatures over plant growth threshold | DAYMET |
|  | Spring mean vapor pressure deficit (AMJ) | VPD | Kpa | mean vapor pressure deficit of spring for observation year | DAYMET |
|  | Total precipitation | PPT | mm | sum of accumulated precipitation for spring of observation year | DAYMET |
|  | Previous mean winter temperature (DJF) | Winter Tmean | °C | mean winter temperature for previous observation winter | DAYMET |

Using a hierarchical Bayesian approach, we modeled DOY of the observed phenological event (leaf-out or flowering) for individual i of species j using a normal likelihood distribution (see Lee et al., 2022):

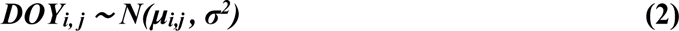

The mean, μ, was modeled with an intercept term (β0), slope terms representing phenological sensitivity to mean spring temperature (β1), elevation (β2), latitude (β3), and species random effects (αj):

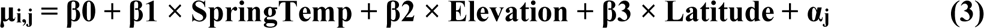

We used slightly informative priors to estimate parameters: β0, β1, β2, β3, αj ∼ N(0, 1E-3); 1/σ2 ∼ Uniform (0,100). Our models used 3,000 burn-in iterations, and three MCMC chains each containing 10,000 iterations. Models were run separately for each stratum for the full AT (i.e., canopy vs. understory, addressing Q1 and Q2), and for each AT region (addressing Q3) combination using the R2jags package (v0.7-1; Su and Yajima, 2022) in R v4.1.0 (R Core Team, 2023). Models for each individual species from both functional groups were also run. Parameter values (means, variances, and covariances) were estimated from posterior distributions and are considered significantly different if the 95% credible intervals (CIs) of their posterior distributions do not overlap. Bayesian R^2^ values were calculated to evaluate variance explained by spring mean temperature on DOY (Gelman et al., 2019). Our parameter values were used to model the direction and magnitude of change to spring phenological windows across the landscape for each AT region in ArcPro v3.1. 1-km rasters of 30-year normal spring mean temperature (DAYMET), elevation (DEM), and latitude were used as raster math inputs to visualize changing phenological windows, but only for areas classified as temperate broadleaf forest using the GAP/LANDFIRE National Terrestrial Ecosystems dataset (USGS, 2016).

For functional group modeling of DOY, we ultimately used 11 of the 14 tree species for spring window calculations. The excluded species were still examined at a species level (see Table 2). We felt this approach was appropriate because these species had relatively few observations (n < 500; e.g., *Sorbus americana*). To test the assumption that understory species flowering time is correlated with leaf-out timing, we re-ran all Bayesian models with DOY_leaf_ of the understory as a response instead of DOY_flower_. General patterns for broad functional groups were not substantially different from one another (Supplemental Table S1).

## Results

### Q1 – Phenological patterns of understory and canopy species

Overall, spring phenology of both canopy trees and understory forest species tended to be earlier when mean spring temperatures (April-June) were warmer across the entire AT Corridor. On average, canopy trees advanced 3.8 days/°C of warming, while understory species as a whole advanced 6.4 days/°C (Table 3). This was true whether looking at understory DOY_flower_ or DOY_leaf_, indicating that comparing understory flowering and canopy leaf-out timing was appropriate as a proxy for understory leaf-out (Supplemental Table S1). Nine of the 11 understory species and six of the 14 canopy tree species significantly (95% CI non-overlapping with zero) advanced their spring phenologies with warmer temperatures (Table 4). For understory species, the spring ephemerals *Dicentra cucullaria* and *Erythronium americanum* displayed the highest temperature sensitivity for flowering (> 6 days/°C), while *Clintonia borealis*, *Viburnum lantanoides*, and *Trientalis borealis* were the least sensitive (< 3 days/°C, Table 4). For canopy tree species with significant sensitivity values, *Fagus grandifolia*, *Acer saccharum*, and *Quercus rubra* showed the most temperature sensitivity for leaf-out (> 4 days/°C), while *Acer rubrum* and *Betula alleghaniensis* were the least sensitive (< 3 days/°C, Table 4). In general, later flowering or leaf-out occurred for individuals at higher elevations or latitudes, consistent with previous studies and predictions under Hopkins’ Bioclimatic law (Table 4).

**Table 3:** Posterior parameter (and 95% credible intervals, CI) estimates of DOY_leaf_ (canopy) and DOY_flower_ (understory) from hierarchical Bayesian modeling for 8 species of understory plants and 11 species of canopy trees across the full AT and partitioned between the three study regions, South (34-38°N), Mid-Atlantic (38-42°N), and North (42-46°N). DIC (deviance information criterion) and Bayesian R^2^ are included. Bolded parameter estimates include 95% CI not overlapping zero.

| Full AT |  |  |  |  |  |  |  |  |  |  |
| --- | --- | --- | --- | --- | --- | --- | --- | --- | --- | --- |
| Region | Functional Group | n | DIC | R<sup>2</sup> | Spring Mean Temperature (°C) | 95% CI | Elevation (m) | 95% CI | Latitude (°) | 95% CI |
| Full AT | Understory | 45293 | 200745 | 0.51 | **-6.12** | -6.43, -5.82 | **30.80** | 28.83, 32.77 | **2.42** | 2.12, 2.72 |
| Full AT | Canopy | 72957 | 657608 | 0.23 | **-3.41** | -3.77, -3.06 | **20.30** | 18.39, 22.31 | **3.87** | 3.58, 4.17 |
| Region-specific |  |  |  |  |  |  |  |  |  |  |
| South | Understory | 10819 | 61861 | 0.30 | **-5.66** | -7.65, -3.63 | **6.25** | 4.29, 8.21 | -3.27 | -5.03, 0.52 |
| South | Canopy | 35242 | 313516 | 0.19 | **-5.86** | -6.49, -5.28 | **13.64** | 11.63, 15.58 | **3.80** | 3.09, 4.58 |
| Mid-Atlantic | Understory | 10834 | 64166 | 0.29 | **-11.45** | -13.56, -9.35 | -0.47 | -2.42, 1.48 | 2.85 | -0.17, 5.83 |
| Mid-Atlantic | Canopy | 13283 | 129899 | 0.17 | **-9.43** | -11.38, -7.46 | 1.98 | -0.09, 3.91 | 0.07 | -2.16, 2.26 |
| North | Understory | 23640 | 179396 | 0.69 | **-6.35** | -6.65, -6.06 | **37.83** | 35.87, 39.82 | **1.64** | 0.95, 2.33 |
| North | Canopy | 24432 | 197718 | 0.41 | **-3.68** | -4.10, -3.25 | **22.15** | 20.22, 24.15 | **3.07** | 2.20, 3.93 |

**Table 4:** Posterior parameter (and 95% credible intervals, CI) estimates of DOY_leaf_ (canopy) and DOY_flower_ (understory) from hierarchical Bayesian modeling for individual canopy tree (gray shading) and understory plant species across the full AT Corridor. DIC (deviance information criterion) is included. Bolded parameter estimates include 95% CI not overlapping zero.

| Species | Common Name | Functional Group | n | DIC | Spring Mean Temperature (°C) | 95% CI | Elevation (m) | 95% CI | Latitude (°) | 95% CI |
| --- | --- | --- | --- | --- | --- | --- | --- | --- | --- | --- |
| Aquilegia canadensis | Red columbine | Understory | 539 | 6398 | **-4.76** | -5.92, -3.54 | 1.17 | -2.61, 4.99 | **3.99** | 3.16, 4.77 |
| Clintonia borealis | Blue-bead lily | Understory | 5816 | 28739 | **-3.10** | -4.06, -2.67 | **26.83** | 24.90, 28.79 | **4.31** | 3.58, 5.09 |
| Cornus canadensis | Canada dogwood | Understory | 5261 | 25607 | **-3.94** | -4.87, -2.87 | **11.96** | 10.03, 13.91 | **4.59** | 2.26, 6.76 |
| Dicentra cucullaria | Dutchman's breeches | Understory | 571 | 7654 | **-6.02** | -7.15, -4.98 | **5.56** | 7.87, 3.33 | **2.89** | 3.65, 2.09 |
| Erythronium americanum | Trout lily | Understory | 5891 | 12382 | **-6.43** | -6.89, -6.07 | **3.10** | 1.13, 5.02 | **2.65** | 1.80, 3.49 |
| Maianthemum canadense | Canada mayflower | Understory | 6975 | 38522 | **-5.06** | -6.34, -4.84 | **12.67** | 10.76, 14.62 | 0.19 | -0.63, 1.01 |
| Sanguinaria canadensis | Bloodroot | Understory | 2434 | 3066 | **-5.27** | -6.79, -4.75 | -0.35 | -2.31, 1.59 | **7.48** | 3.11, 11.74 |
| Trientalis borealis | Starflower | Understory | 535 | 2217 | -0.42 | -3.20, 2.67 | **3.42** | 1.45, 5.38 | **-5.47** | -8.78, -2.20 |
| Trillium erectum | Red trillium | Understory | 6018 | 29870 | **-5.74** | -7.94, -6.39 | **4.92** | 2.98, 6.85 | **1.17** | 0.44, 1.89 |
| Trillium undulatum | Painted trillium | Understory | 6152 | 22471 | **-4.36** | -5.84, -3.42 | **14.33** | 12.38, 16.23 | **3.47** | 2.81, 4.13 |
| Viburnum lantanoides | Hobblebush | Understory | 5101 | 32503 | **-3.53** | -4.68, -2.82 | **17.27** | 15.26, 19.28 | **7.63** | 6.46, 8.85 |
| Acer pensylvanicum | Striped maple | Canopy | 7901 | 63799 | **-3.72** | -4.95, -2.49 | **12.01** | 10.00, 14.00 | **3.55** | 2.50, 4.59 |
| Acer rubrum | Red maple | Canopy | 26926 | 241645 | **-1.71** | -2.25, -0.96 | **10.50** | 8.58, 12.54 | **4.86** | 4.33, 5.42 |
| Acer saccharum | Sugar maple | Canopy | 9422 | 84363 | **-4.02** | -5.03, -3.10 | **-2.34** | -4.24, -0.31 | **2.62** | 1.85, 3.47 |
| Betula alleghaniensis | Yellow birch | Canopy | 6069 | 49747 | **-2.61** | -3.24, -1.47 | **14.53** | 12.63, 16.53 | **4.66** | 3.86, 5.48 |
| Betula lenta | Sweet birch | Canopy | 3548 | 32373 | 0.56 | -0.94, 2.35 | **12.67** | 10.61, 14.58 | **7.71** | 6.68, 8.76 |
| Betula papyifera* | Paper birch | Canopy | 270 | 2347 | -0.03 | -6.12, 6.08 | **2.11** | 0.14, 4.02 | -4.79 | -11.69, 3.83 |
| Fagus grandifolia | American beech | Canopy | 8528 | 68346 | **-5.78** | -6.36, -4.35 | **2.22** | 0.33, 4.14 | 0.74 | -0.04, 1.52 |
| Fraxinus americana | White ash | Canopy | 790 | 7506 | 0.45 | -3.37, 4.36 | 1.86 | -0.09, 3.80 | **5.08** | 0.13, 10.13 |
| Ostrya virginiana | Ironwood | Canopy | 674 | 6324 | -0.54 | -4.40, 2.48 | **7.60** | 5.65, 9.62 | **9.42** | 5.25, 13.46 |
| Prunus pensylvanica* | Pin cherry | Canopy | 360 | 3208 | -1.95 | -9.82, 6.50 | -1.69 | -3.62, 0.31 | -6.92 | -22.07, 8.25 |
| Prunus serotina | Black cherry | Canopy | 2966 | 27533 | 0.80 | -0.42, 2.13 | **6.41** | 4.51, 8.37 | **8.39** | 6.45, 10.32 |
| Quercus rubra | Red oak | Canopy | 4035 | 36833 | **-4.71** | -6.34, -2.40 | -0.69 | -2.63, 1.26 | 0.15 | -1.14, 1.43 |
| Sorbus americana* | American mountain-ash | Canopy | 418 | 3604 | -2.31 | -12.47, 8.96 | **-5.37** | -7.30, -3.41 | -5.88 | -27.30, 17.22 |
| Tilia americana | American basswood | Canopy | 1050 | 9845 | -0.90 | -3.48, 1.32 | **2.00** | 0.04, 3.95 | **5.39** | 2.25, 8.58 |

\*Species was modeled independently and not used in broad functional group modeling.

### Q2 – Differences between functional groups

Spring-flowering forest understory species responded more strongly to warmer temperatures than did canopy trees when data were pooled across the entire length of the AT Corridor – by approximately 1.6 days for every 1 °C increase in spring mean temperature (Table 3). In particular, the understory species *Dicentra cucullaria*, *Erythronium americanum, Sanguinaria canadensis*, and our two *Trillium* species were much more sensitive to potential temperature increases than most individual canopy tree species, with the exception of *Fagus grandifolia* (Table 4). Greater understory sensitivity to temperature than canopy trees suggests an expansion of the spring phenological window. Our models revealed that greater window expansion was likely to occur at higher elevations and latitudes (Table 3).

### Q3 – Latitudinal patterns of phenology

For southern and mid-Atlantic regions (34–38°N and 38–42°N, respectively), there were no detectable differences between functional groups in responsiveness of spring phenology to temperature (overlapping 95% CI). At northern latitudes (42–46°N), understory species as a group advanced their spring phenologies more strongly than trees with respect to temperature, by around 2.6 days/°C (Table 3, Figure 4). Both functional groups advanced their phenologies nearly twice as much in the mid-Atlantic region (38-42°N) than either southern or northern AT regions (Figure 4). Taken together, the phenological window likely experienced expansion to a greater degree in northern latitudes, and in general, at higher elevations for all regions examined (Figure 5). Spring windows expanded little or remained stable at mid-to southern latitudes (Figure 5). We found that annual mean spring temperatures in our dataset increased for all regions over the timespan analyzed (2004-2022), but only significantly at higher latitudes (Figure 6), indicating greater warming at northern latitudes and more variation in temperature for which plants can respond.

**Figure 4:**
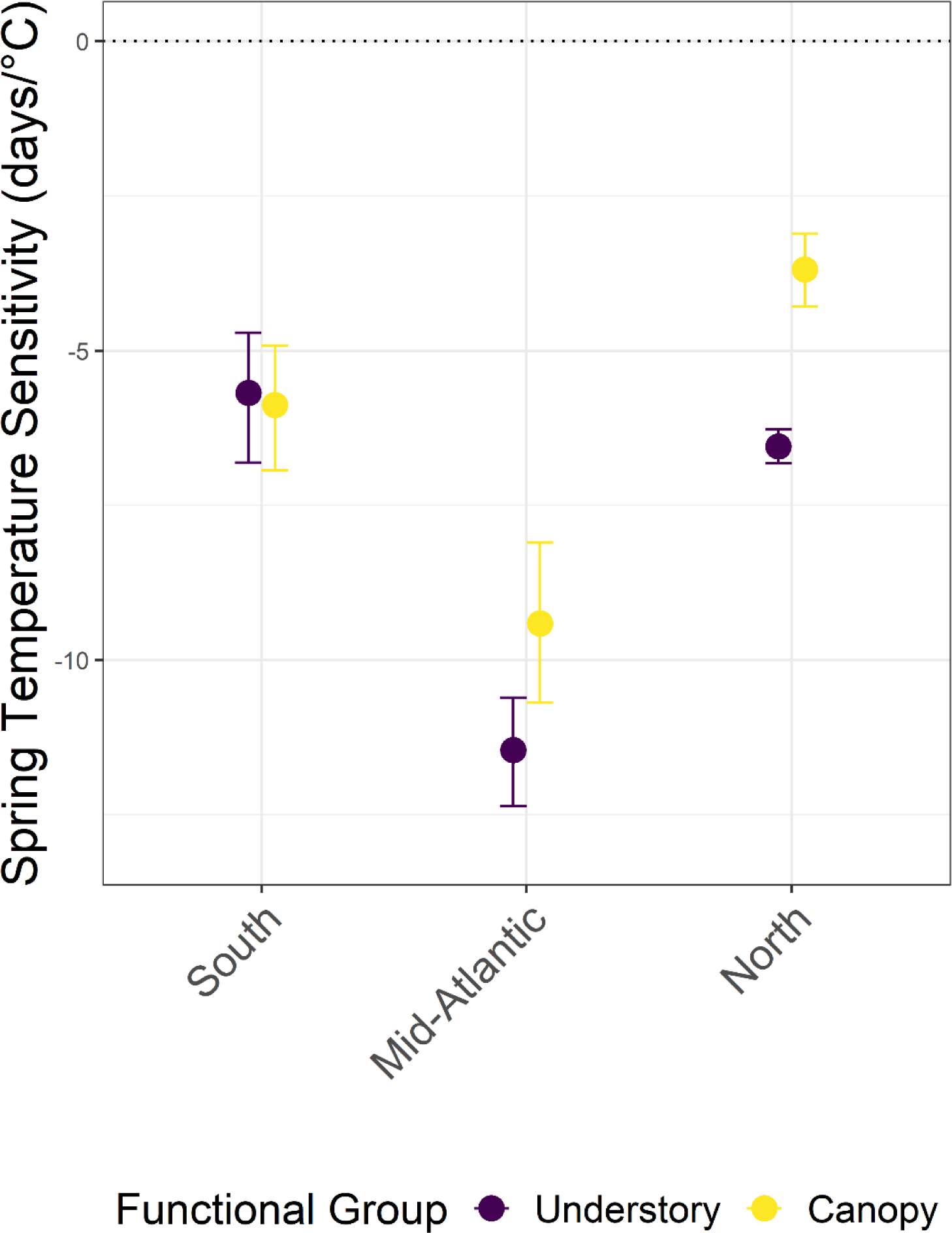
Spring temperature sensitivity (number of days of phenology advance per 1°C of warming) for both canopy trees and understory plants (as a whole), partitioned between our three study regions (with 95% CI). Negative y-axis values indicate earlier phenology with warming. The Northern region displays evidence for phenological window expansion, with a stable window illustrated for the Southern region. Understory species also display greater temperature sensitivity than canopy trees in the Mid-Atlantic region, although 95% CI overlap for both functional groups.

**Figure 5:**
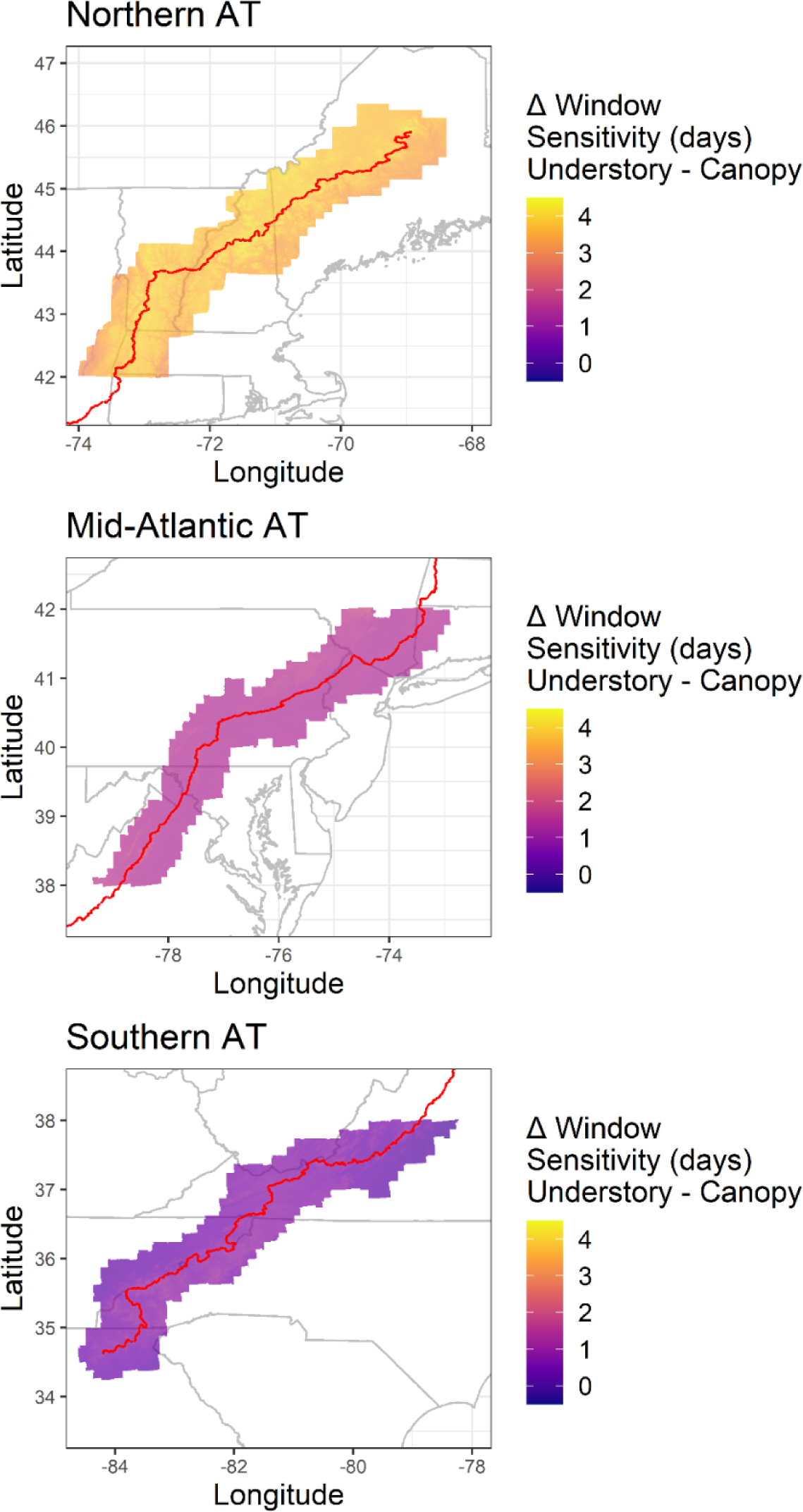
Changes (Δ) in spring phenological windows between understory plants and canopy trees, as influenced by temperature, elevation and latitude, in each study region along the AT Corridor. Warmer colors indicate greater possible phenological window expansions whereas cooler colors show smaller expansions, no differences, or contractions of the spring window. Greater window expansions occur more commonly at higher elevations and latitudes, particularly in the Northern region. Changes are more subtle in the Mid-Atlantic and Southern regions. The red line indicates the AT.

**Figure 6:**
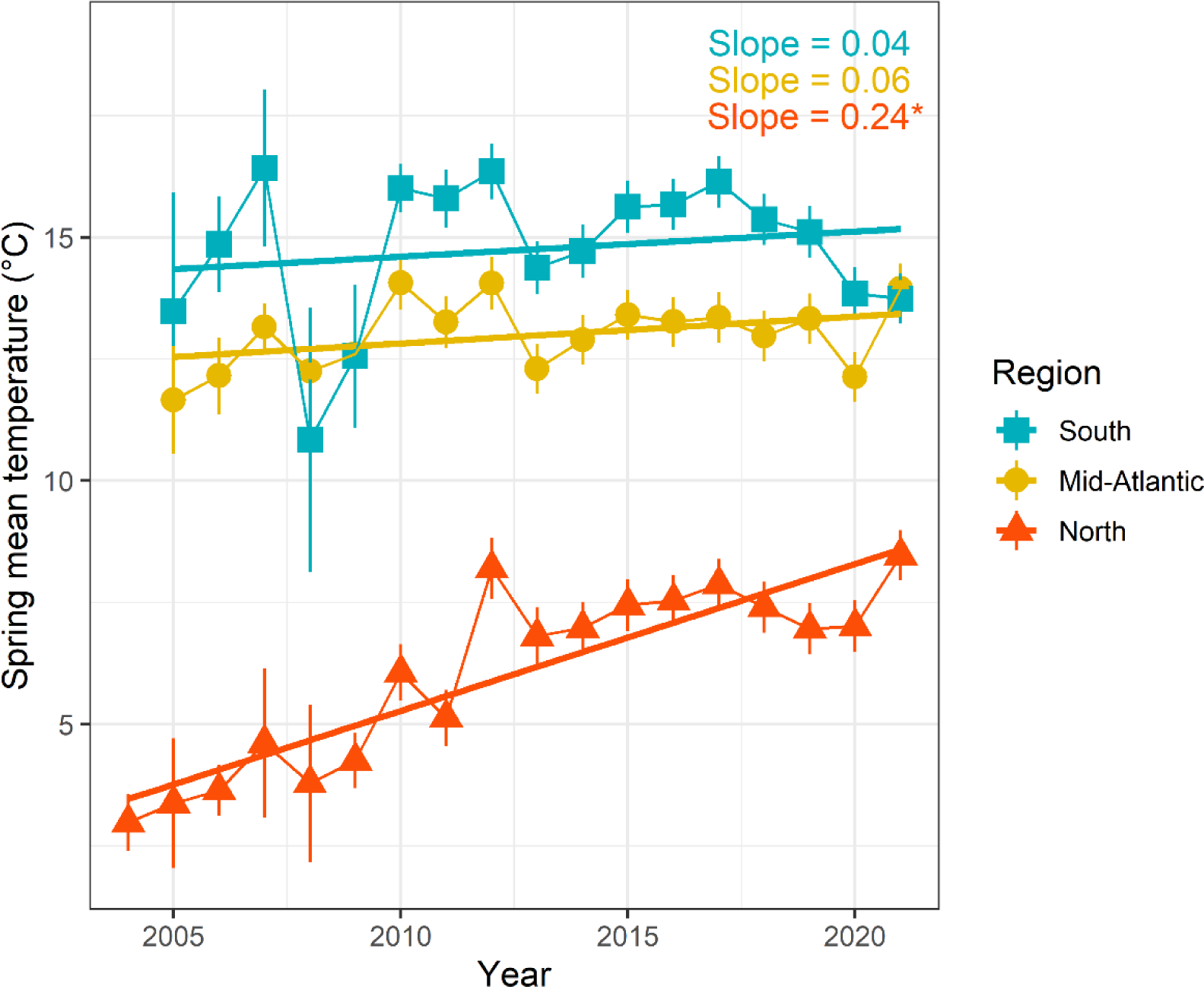
Annual spring (April-June) mean (±SE) temperatures between 2004-2022 across the AT Corridor, partitioned between three latitudinal bands (South: 34-38°N, blue squares; Mid-Atlantic: 38-42°N, orange circles; North: 42-46°N, red triangles). Linear fits display trends of increasing temperature through time, although only the Northern region experienced significant warming (see top-right for coefficient estimates and significance; *p<0.05). Temperatures were estimated from DAYMET daily measurements from all observation locations. Note that annual spring mean temperatures were not recorded for the South and Mid-Atlantic regions in 2004 due to poor data quality.

## Discussion

Overall, we find regional disparities of plant spring phenological response to warming for species within temperate deciduous forests of the eastern United States. Specifically, we illustrate that spring phenology is advancing with temperature for both spring blooming forest understory species and canopy trees in North America’s eastern hardwood forest ecosystems (addressing Q1). Spring phenology is also advancing to a greater degree at both higher latitudes and elevations for most species examined. However, understory species’ phenologies are advancing at a greater rate than canopy tree leaf-out phenologies, but only significantly so at northern latitudes (42-46°N) with no detectable difference between functional groups for lower and middle latitudes (34–42°N, addressing Q2). Furthermore, both functional groups vary in their phenological response to warmer temperatures across the latitudinal range, with understory plants and canopy trees at middle latitudes advancing their spring phenologies more than those at lower and higher latitudes (addressing Q3). Given that both functional groups were more sensitive to temperature at middle latitudes (38–42°N) compared to low (34–38°N) and high latitudes (42–46°N), other unmeasured factors may be at play influencing the temporal dynamics of the spring phenological window. Since a high proportion of mid-Atlantic observations were taken in urban areas (many near the New York City metropolitan area), one unexplored possibility is that the temperature sensitivity of this region may be due to effects from land-use (Luo et al., 2007; Zipper et al., 2016).

The phenological advance of both deciduous tree and spring-flowering understory plants is supported by previous studies and suggests that rising spring temperatures could increase the length of the growing season in temperate deciduous forests (Monahan et al. 2016; Melaas et al., 2018; Seyednasrollah et al., 2020; Moon et al., 2021; Li et al., 2022). However, while our finding that understory spring phenology is advancing faster than deciduous canopy trees under warmer temperatures at northern latitudes (42–46°N) agrees with a recent study (Alecrim et al., 2022), it is at odds with other studies conducted within the same region (Heberling et al., 2019; Lee et al., 2022). The variation in canopy-understory phenology results across these studies may reflect differences in methodological approaches, including the density of observations, and the study species (see Alecrim et al., 2022; see below for possible explanations).

The observed differences in temperature sensitivity and potential changes to the spring phenological window could be attributed to either disparities in changing environmental conditions or differences in forest community composition across our latitudinal gradient. Calculated mean spring temperatures (April-June) reveal warming trends for all regions examined, however, only the northern AT experienced statistically significant warming between 2004-2022 (Figure 6). While the Mid-Atlantic region displayed greater temperature sensitivities for both understory plants and canopy trees than other regions, only in the northern AT did we find significantly different sensitivities between the two functional groups. It is possible that consequences of greater warming in the north are altering plant responses to a changing climate in ways at odds with conspecific southern populations (see below). Of course, disparate community composition which encapsulate our study species across regions may also be driving these patterns.

The northern AT region differs from lower latitudes in several other relevant ways. Most notably, the northern AT is characterized by longer and colder winters, deeper snowpack, shorter growing seasons, and is projected to warm faster than other regions over the course of this century (US National Climate Assessment, 2018 (https://nca2018.globalchange.gov/), Janowiak et al., 2018). These trends may be relevant in several ways. First, chilling requirements for understory plants may still be met in the north despite recent warming (at least up to a certain threshold not yet reached), meaning that these species will not suffer reduced performance and could, at least in the short term, benefit from a longer spring window before canopy closure (Zhang et al., 2007 Prevéy et al., 2017). Second, a decreasing snowpack, particularly in the north, would decrease soil temperatures in the late winter period (Zhu et al., 2019; Zohner et al., 2017), but would also allow for a greater time for herbaceous species to be uncovered by snow in the spring, a critical time for growth (Marchin et al., 2015; Augspurger and Salk, 2017; Contosta et al., 2017). While during the summer months understory temperatures are buffered (cooler) by the canopy, this is not the case prior to canopy closure in the spring – suggesting that understory species could be more directly influenced by surface air temperatures than has been previously suggested (Richardson and O’Keefe; De Frenne et al., 2011; Jacques et al., 2015; De Frenne et al., 2021). To resolve these various interacting factors, future work must measure relevant climate covariates, such as snow cover and soil temperature, in locations that record phenology of individual plants.

The temporal dynamics of spring phenology are hard to predict given the high variation in published sensitivities (e.g., Heberling et al., 2019; Alecrim et al., 2022; Lee et al., 2022). We argue that methodological differences between these disparate studies may be the root cause of the observed discrepancies. First, the phenophase used to model functional group temperature sensitivity (leaf-out vs. flowering) can influence the interpretation of the spring phenological window. While we found that leaf-out and flowering timing were correlated in our study (and see Heberling et al., 2019), appropriate ancillary data is required to successfully utilize each metric. Namely, some kind of intensity value is needed to describe a distribution of these events, rather than a presence/absence record (Buonaiuto et al., 2021). Without this information, it is difficult to know the exact time of flowering or leaf-out. Second, a substantial amount of variation in both temporal and spatial ranges examined could lead to the observed differences in reported results. It would be difficult to compare the results of two phenological window studies examining advancing phenology with an order of magnitude difference in the time record illustrated (decadal vs. century timeframes), as the magnitude of warming is dissimilar (Ge et al., 2015; Alecrim et al., 2022). Likewise, results from studies examining specimens at a local scale may not be applicable to a regional-scale given the exponential growth of environmental variation encountered, especially in the context of microclimates and climate refugia (Wielgolaski, 1999; Wolkovich et al., 2021; Pastore et al., 2022). Third, the size and source of the dataset used may be critical for phenological studies. Large phenology datasets, as is the case with our study, are preferable to smaller ones for regional-scale studies; however, access to such rich data sources are not always possible.

This study is unique in this area given our large sample size and use of citizen-science derived iNaturalist data, which served to greatly expand the temporal and spatial variation described in our study region (Supplemental Figure S1). We recommend that similar future research engage with this efficacious resource. Further, while spring ephemerals and other herbaceous species have been the focus of understory phenology patterns, tree seedlings have largely been ignored in their phenological response to changing climate (Augspurger and Bartlett, 2003; Lopez et al., 2008; Richardson and O’Keefe, 2009; but see Lee and Ibanez, 2021a). Seedlings represent the future composition of a forest, and any change in seedling survival and growth related to shifting phenology is important to capture (Lee and Ibanez, 2021a; 2021b). We advocate for a stronger emphasis on observations of woody seedling species in the understory moving forward.

Earlier flowering and leaf-out relative to canopy closure could serve to benefit plant performance of understory forbs and shrubs through a number of mechanisms. Advanced leaf-out and flowering could trigger an increase in photosynthate accumulation and storage (Keenan et al., 2014; Teets et al., 2023). Greater access to resources could also provide a boost to both vegetative growth and reproduction (Kudo et al., 2008; Heberling et al., 2019). Thus, assuming understory plant fitness is not affected by other changes caused by shifting phenology or other climate changes such as extreme precipitation or drought, an expansion of the phenological window could make these species more resilient to a changing climate. This is especially true for spring ephemeral species which almost entirely rely on high light availability in the spring. Indeed, our results reveal that the most temperature sensitive species were ephemeral species such as *Dicentra, Erythronium, Trillium,* and *Sanguinaria*.

Advancing understory spring leaf-out and flowering may also have negative impacts on plant performance. First, earlier leaf-out could expose both spring ephemerals and trees to unpredictable late frost (or “false spring”) events which can cause significant physical damage and loss of fitness, particularly as climate change makes extreme frost events more common (Augspurger, 2009; Marino et al., 2011, Casson et al. 2019). Leaf or leaf-bud loss to frost represents a significant cost for deciduous trees, affecting growth, reproduction, canopy expansion and nutrient reserves, as refoliation to compensate for damage demands extra resources (Inouye, 2008; Augspurger, 2009; Pardee et al., 2019). Second, plant phenological shifts relative to herbivores and pollinators may have a large effect on plant performance. For instance, shifts in timing of herbivore emergence relative to plant phenology, as well as changes in the frequency or severity of herbivore outbreaks could have major impacts on understory shading and carbon budgets of trees and understory plants (Kudo et al., 2008; Weed et al., 2013). Importantly, if plants and insects do not respond at the same rate to warming, mismatches between flowers and flower-visitors could occur (Kudo and Ida, 2013; Petanidou et al., 2014; Forrest, 2015; Kudo and Cooper, 2019). While not as relevant for wind-pollinated deciduous trees, forest understory forbs are generally insect-pollinated, and given the short flowering period of these species, phenological mismatches between these plants and their pollinators are possible (Kudo and Ida, 2013; Kudo and Cooper, 2019).

### Conclusions

Here, we find evidence that understory plants in eastern North America are advancing their spring phenologies 1.6 days/°C faster than canopy trees near full leaf-out; in other words, this functional group appears more sensitive to air temperature increases than trees. The expansion of the spring phenological window could be a net positive for understory plant performance under changing climate conditions; however, many other unexplored phenomena, such as biotic interactions and climate-induced hydrologic variability, make forecasting changes to forest communities challenging. We note distinct patterns of phenological sensitivity across a latitudinal gradient, indicating that forest plant response to warming in eastern forests will not be uniform across space. It is also difficult to determine how specific species will ultimately respond to warming, or how long-term phenological dynamics will be altered (i.e., do threshold responses to warming exist for these species?). We posit that more spatially diverse phenological data, particularly from citizen-science driven efforts, can help inform research related to forest resilience to climate change, and that future work would greatly benefit from a more standardized approach.

## Acknowledgements

We thank the National Phenology Network, the US Forest Service, the US National Park Service, Appalachian Mountain Club staff, interns and volunteers and citizen scientists across the AT Corridor. Doug Weihrauch initially helped conceive AMC’s Mountain Watch Project. We thank Dr. Caitlin McDonough MacKenzie for her feedback on earlier versions of this manuscript. We also thank Dr. Ben Lee for helpful comments on methods and code used to perform our analyses, as well as Miriam Ritchie for assistance with some figures. Funding was provided by the AMC’s Ken Kimball Research Fellowship, the Waterman Fund, the Forest Ecosystem Monitoring Cooperative (FEMC), a National Geographic Explorer Grant, the Adelard A. and Valeda Lea Roy Foundation, and the Appalachian Trail Conservancy’s (ATC) Wild East Action Fund. Some data were provided by the USA National Phenology Network and the many participants who contribute to its Nature’s Notebook program. Some data were also accessed through the iNaturalist platform and its many volunteer observers.

## Author Contributions

JT conceived the research questions with input from GM and SN. JT planned and designed the research and conducted all data analysis with input from GM and SN. JT wrote the manuscript, and all authors contributed substantial revisions and edits.

