## Supplemental for "Distinct latitudinal patterns of shifting spring phenology across the Appalachian Trail Corridor"

**Supplemental Information**

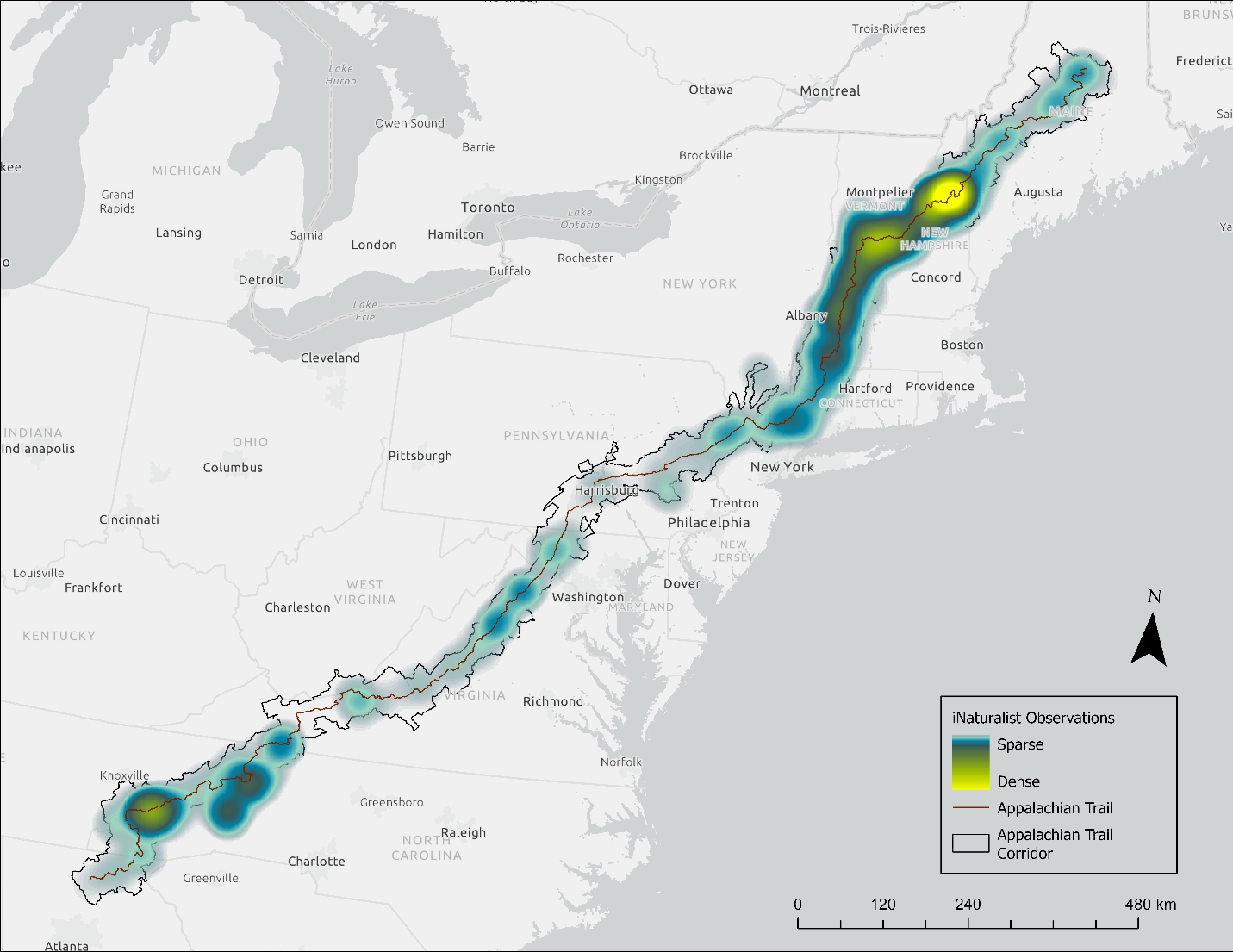

**Supplemental Figure S1:** Visualization of the density of iNaturalist observations for both canopy trees and woodland understory species between 2018 and 2022 (n ~ 40,000) across the entire length of the AT Corridor.

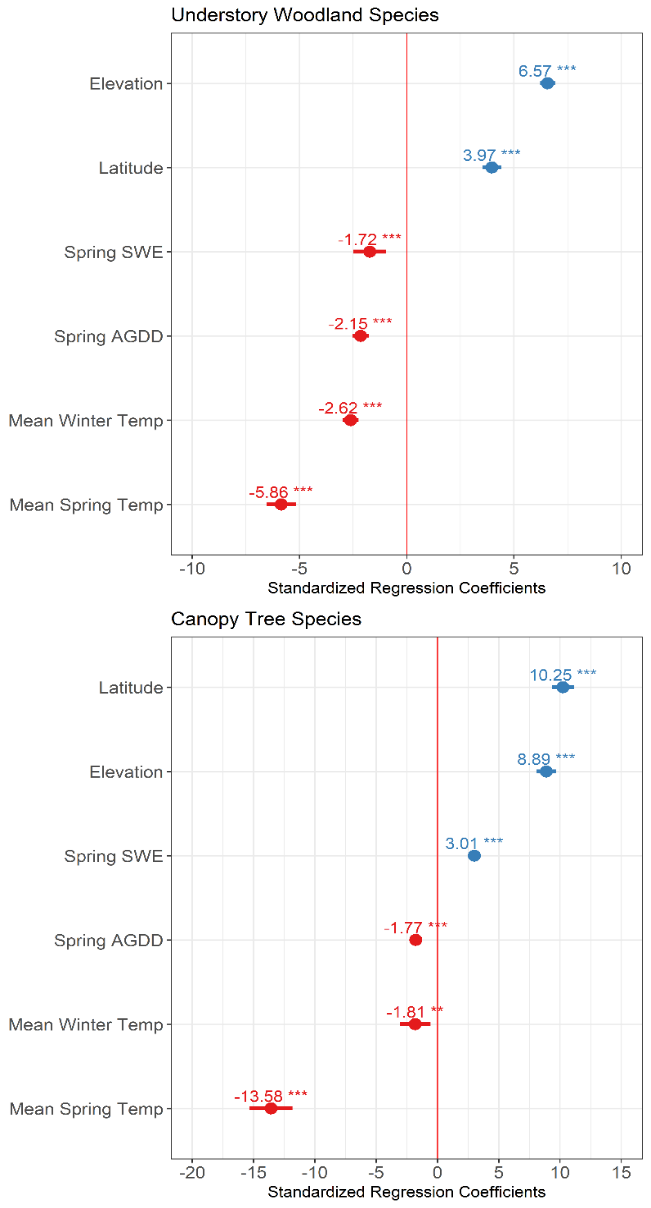

**Supplemental Figure S2:** Coefficient estimates of candidate climate and landscape variables from initial exploratory regressions. All variables were scaled and centered prior to model initialization. DOY of leaf-out (canopy) or flowering and leaf-out (understory) was used as a response for all models. Spring (April-June) mean temperature, latitude, and elevation consistently had significant effects on DOY for both functional groups. Abbreviations and units: Latitude (°N), Elevation (m a.s.l.), Spring SWE (snow water equivalent, mm), Spring AGDD (accumulated growing degree days), Mean Winter Temp (°C), Mean Spring Temp (°C).

**Supplemental Table S1:** Posterior parameter (and 95% credible intervals, CI) estimates from hierarchical Bayesian modeling for 11 species of understory plants across the full AT and partitioned between the three study regions, South (34-38°N), Mid-Atlantic (38-42°N), and North (42-46°N). Response is DOY_leaf_ for all understory species to facilitate comparison with DOY_flower_ (from Table 1). DIC (deviance information criterion) and Bayesian R^2^ are included. Bolded parameter estimates include 95% CI not overlapping zero.

| **Full AT** |  |  |  |  |  |  |  |  |  |  |
| --- | --- | --- | --- | --- | --- | --- | --- | --- | --- | --- |
| **Region** | **DOY (leaf or flower)** | **n** | **DIC** | **R^2^** | **Spring Mean Temperature (°C)** | **95% CI** | **Elevation (m)** | **95% CI** | **Latitude (°)** | **95% CI** |
| Full AT | Leaf | 16991 | 152100 | 0.41 | **-5.04** | -6.40, -3.71 | **22.95** | 21.00, 24.99 | **2.29** | 2.08, 2.52 |
| Full AT | Flower | 45293 | 200745 | 0.51 | **-6.12** | -6.43, -5.82 | **30.80** | 28.83, 32.77 | **2.42** | 2.12, 2.72 |
| **Region-specific** |  |  |  |  |  |  |  |  |  |  |
| South | Leaf | 920 | 8791 | 0.28 | **-5.51** | -7.45, -3.53 | **5.15** | 4.40, 5.89 | **2.36** | 0.37, 4.35 |
| South | Flower | 10819 | 61861 | 0.30 | **-5.66** | -7.65, -3.63 | **6.25** | 4.29, 8.21 | -3.27 | -5.03, 0.52 |
| Mid-Atlantic | Leaf | 1334 | 14165 | 0.27 | **-10.53** | -12.46, -8.59 | 1.38 | -0.07, 2.73 | -1.86 | -3.79, 0.11 |
| Mid-Atlantic | Flower | 10834 | 64166 | 0.29 | **-11.45** | -13.56, -9.35 | -0.47 | -2.42, 1.48 | 2.85 | -0.17, 5.83 |
| North | Leaf | 14737 | 137334 | 0.55 | **-5.88** | -6.06, -3.70 | **22.15** | 20.22, 24.15 | **1.94** | 1.32, 3.26 |
| North | Flower | 23640 | 179396 | 0.69 | **-6.35** | -6.65, -6.06 | **37.83** | 35.87, 39.82 | **1.64** | 0.95, 2.33 |
